# Drivers of Variation of the Zebrafish Egg Microbiome

**DOI:** 10.1101/2025.09.01.673521

**Authors:** Caitlin Smith, Karen Adair, Brendan Bohannan

## Abstract

Microbiomes are integral to the fitness of many animals, yet little is known about early-life microbiome assembly. What is known largely comes from studies in non-model organisms, where it is difficult to distinguish host genetics from the environment as drivers of microbiome assembly.

Here, we used a popular vertebrate model, zebrafish (*Danio rerio*), to address the major drivers of environmental assembly in 124 individuals. We surveyed the egg microbiome from fertilization to hatching using 16S rRNA gene sequencing and investigated development stage and parentage as potential drivers of microbiome variation. Phylogenetic diversity of the egg microbiome decreased during development, suggesting that environmental filtering may be a major driver of community assembly. Parentage also had an influence on microbiome assembly; eggs derived from the same clutch were often more similar in microbiome composition compared to eggs derived from different clutches. The effect of parentage on microbiome similarity decreased over development, suggesting that parents may be an initial source of egg microbiome members.

The egg microbiome undergoes dramatic compositional shifts during development. Understanding the drivers of this variation is important to consider, especially given the role of the microbiome in health and development, and that early exposure to microbes may shape later animal development. Our findings suggest that parents may serve as an important source of microbial symbionts to their offspring, even in an animal that does not provide parental care, such as laboratory zebrafish, and that the developing egg may play an increasingly important role in driving microbiome variation across individuals.

**Importance:** Early-life microbiome composition is integral for establishing the normal functioning of multiple host systems, including the nervous system, gastro-intestinal tract, and immune system. However, the drivers of variation in early-life microbiome composition are underexplored, especially in *Danio rerio* (zebrafish), an important model organism that has allowed extensive exploration of the role of the microbiome in host health and development. Here, we characterize the microbiome of individual zebrafish eggs and determine the effects of parental identity and developmental time on shaping the egg microbiome prior to hatching.

Both parentage and age influence microbiome composition and diversity. We found evidence that species-sorting (i.e. selection of microbes by the egg) is an important driver of microbiome assembly during egg development. Furthermore, the effect of parentage on the microbiome, while significant throughout embryonic development, decays over time. Despite the lack of parental care in laboratory reared zebrafish, these findings suggest that the parents are significant source of microbes to their offspring.

## Introduction

The host-associated microbiome is integral to the health and development of animals, especially vertebrates. The gut microbiome is especially important in early life, influencing the development and function of the immune system, nervous system, and GI tract (1–9). Given the importance of the gut microbiome to the development of young vertebrates, determining the mechanisms behind early-life assembly of the microbiome is crucial for a complete understanding of the interactions between vertebrates and their resident microbes. While the importance of the early-life microbiome has been described in the studies above, relatively little is known about the assembly of microbiomes early in life. Those few studies that address early-life microbiome assembly focus on humans (10,11), livestock (12,13), some captive and wild birds (14–18), and invertebrates (19–21). Interpretations of early-life microbiome studies in humans, wild animals, and non-model organisms can be impeded by small sample sizes, lack of environmental control, and long development time. These limitations highlight the importance of understanding the mechanisms of early-life assembly in model vertebrates, such as zebrafish and mice, reared in controlled environments.

Zebrafish are a powerful model organism with which to understand host-microbe interactions and their impact on host health. They are useful for studying how microbiomes assemble, particularly in the gut. Such studies have revealed the processes that drive variation in the gut microbiome across individuals and through development (22–26). However, most studies to date have focused on microbiome assembly in larva and adults, rather than in earlier life stages, such as the egg. Although little is known about early-life microbiome assembly in the zebrafish, it has been shown that interactions between the zebrafish and its gut microbiome early in life influence many aspects of zebrafish health and functioning. For example, experiments on axenic (germ-free), gnotobiotic, and conventionalized or conventionally raised zebrafish larvae have revealed the important roles that specific members of the zebrafish microbiome play in proper neurodevelopment (3,6,27,28), the onset of social behavior (3,28), immune system development (7) and intestinal development (1,2). The sources of these beneficial microbes are not yet well understood.

Eggs are an especially underexplored life stage in zebrafish microbiome development. The egg microbiome may influence health and survival of early-stage fish, as well as the microbiome composition and function in later developmental stages (29). For instance, the microbiome composition of Atlantic salmon eggs can influence the gut and skin microbiome composition of the larvae (30). In zebrafish, axenic derivation of eggs and re-conventionalization of larvae can lead to different microbiome compositions than in conventionally raised larvae (27). This study aims to fill this knowledge gap by determining the drivers of variation in the zebrafish egg microbiome.

A possible influence on microbiome composition of eggs is parental identity, or clutch identity. These differences could be caused through genetic differences (31) or transmission of microbes. For instance, parental microbiome transmission is an important driver of early life assembly in other teleost fish, such as pipefish (*Sygnathus typhle*; (32) and discus fish (*Symphysodon aequifasciata*; (33), but its role in zebrafish is not clear. In laboratory aquaria, zebrafish reproduce via broadcast spawning and are not known to provide parental care (although little is known about the parental care behaviors of wild populations beyond oviposition; (34,35)). Transmission may occur between parent and offspring during reproduction (as has been shown for other fish species; (32) and through parental care (33), although to our knowledge, this has not been previously documented in zebrafish. Parents may also influence the egg microbiome through maternally transmitted immune proteins (36).

In addition to possible parental influences, the assembly of the egg microbiome may also be influenced by changes to the egg as it develops. This has been observed in other fish species, such as Atlantic salmon (29), whitefish (37) and lake sturgeon (38). The developing embryo may also introduce selection pressure on its microbes, as anti-microbial compounds are produced by the embryo prior to hatching (39).

In this study, we address whether clutch identity (i.e. parentage) and development influence the microbiome of zebrafish eggs from 0 to 2 days post fertilization (dpf). To investigate the roles of parentage and development, we crossed wildtype adult zebrafish and collected fertilized embryos for microbiome characterization with 16S rRNA gene sequencing. Specifically, we address the following questions: 1) does microbial diversity change over the course of embryonic development? 2) do clutch identity or developmental stage explain variation in microbiome composition between eggs? and 3) does the effect of shared parentage on microbiome similarity shift over development? For simplicity, developmental stage and age are used interchangeably to refer to the numbers of days post fertilization of the eggs.

## Methods

### Ethics Statement

All zebrafish experiments were approved by the University of Oregon Institutional Animal Care and Use Committee (AUP 20-16 and AUP-21-31).

### Sample Collection

We bred adult wildtype zebrafish (ABCxTu genotype) according to the protocols for pairwise breeding of natural crosses in false bottom containers as detailed in *The Zebrafish Book* (40) and as described in the supplemental methods. Briefly, 20 male/female pairs from two stock tanks (tank “A” and “B”; Figure 1) were placed in crossing tanks with false-bottom inserts to separate fertilized eggs from adults. Eggs were collected and rinsed in embryo media the following day using a tea strainer. Clutches of eggs were further divided across multiple petri dishes and kept at densities of approximately 75 egg per dish. The first clutch from tank A was rinsed and sorted starting at approximately 1 hour post fertilization (hpf), with the last clutch from tank B rinsed and sorted at approximately 8 hpf. Eggs were collected for DNA extraction at 0 days post fertilization (dpf), 1 dpf and 2 dpf.

**Figure 1:**
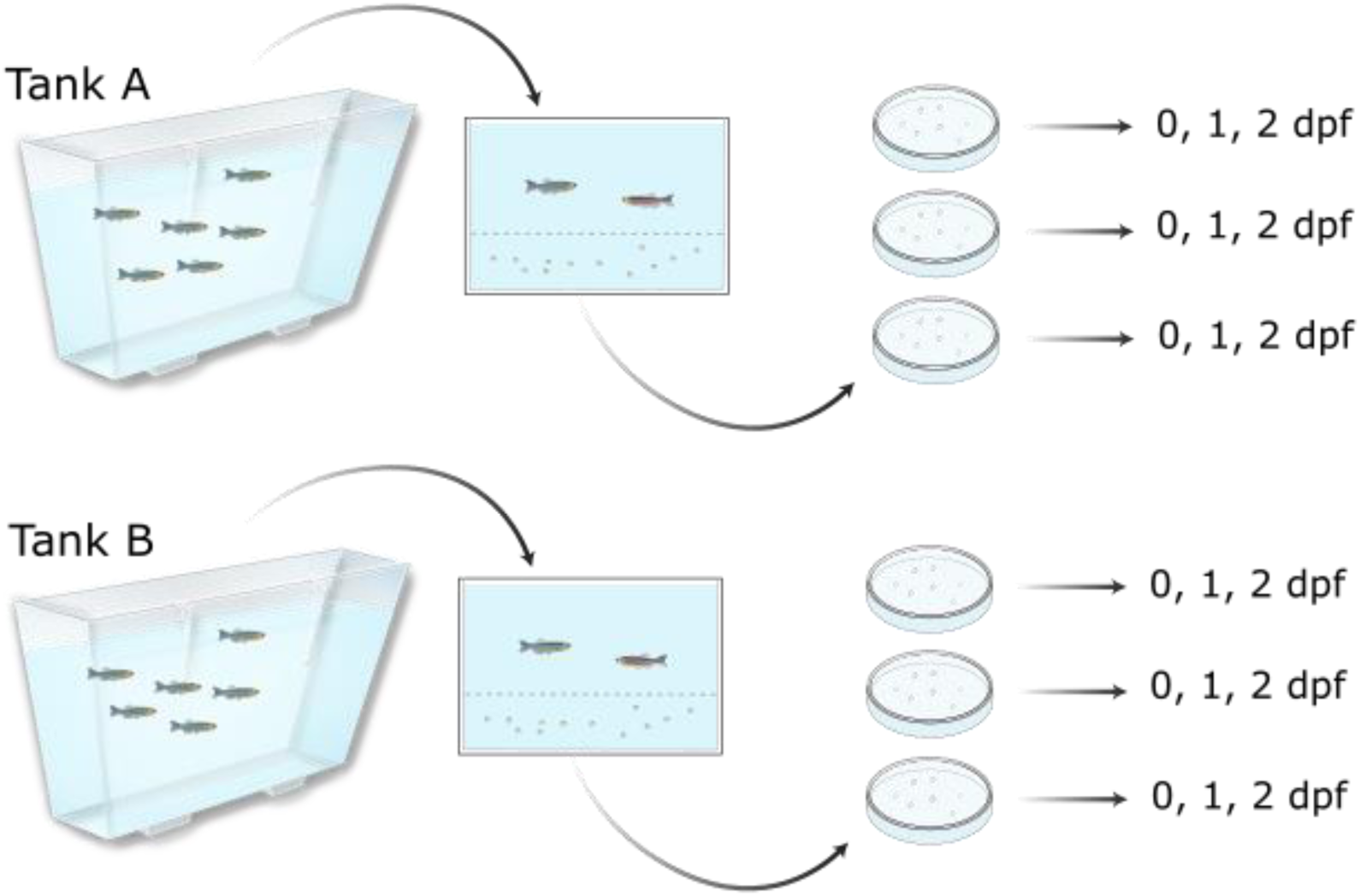
Sampling schematic. Adults were raised in two stock tanks (A and B). Male/female pairs were placed in breeding tanks overnight. Eggs deriving from each successful pair were rinsed and stored in Petri dishes at a density of 50-75 eggs/dish. Five eggs were collected from each dish at 0 dpf, 1 dpf, and 2 dpf.

### Data Analysis

#### Diversity metrics

All data analysis following sequence processing was performed in R version 4.3.2 (41), with *ggplot2* (42), *cowplot* (*43*), and *microshades* (44) used to produce plots. To minimize the effect of library size on variation in alpha and beta diversity, we normalized ASV counts through repeat rarefaction (45,46) and used the *vegan* package to rarefy the ASV table without replacement to a minimum sampling depth of 2,651 reads. We then calculated alpha diversity metrics on the rarefied ASV table using the packages *vegan* (ASV richness, Shannon-Weaver Index/Shannon (47), Pielou’s Evenness Index (48) and *picante* (Faith’s Phylogenetic Diversity, (49,50). For beta diversity, we calculated weighted and unweighted Unifrac (51) and Jaccard and Bray-Curtis dissimilarity (52,53) using *phyloseq* and *vegan*. We repeated this rarefaction step and recalculated diversity 10,000 times using the *parallel* package and averaged the alpha and beta diversity metrics across each randomly rarefied community.

#### Influence of tank, clutch and age on microbiome diversity

Regression models using the package *lmerTest* (54) were used to analyze the effect of development and clutch on mean bacterial 16S copies per egg and alpha diversity. We ran separate models to test how alpha diversity differed by age and between clutches, where sampling depth was provided as a covariate in every test. For regression models exploring the effect of age on alpha diversity, we allowed the intercept to vary between clutches, and when exploring the effect of clutch identity on alpha diversity, we allowed the intercept to vary between ages. We used Tukey tests to conduct multiple comparison tests of mean alpha diversity between groups (i.e. between different ages and between different clutches) with the *multcomp* package (55). P-values for these comparisons were adjusted using the Holm method.

To investigate phylogenetic clustering or overdispersion in the microbiome, we calculated the Net Relatedness Index (NRI; (56,57)) and the Nearest Taxon Index (NTI; (56,57)) of each egg microbiome using the package *picante*, calculating mean pairwise distance (MPD) and mean nearest taxon distance (MNTD) and subtracting 1 from the z-score. We tested whether the mean NRI and NTI were significantly different from zero using a Wilcoxon Signed Rank test.

To test for the effect of clutch on microbiome composition while accounting for variation among Petri dishes (similar to a tank effect observed in cohoused animals; (58,59), we performed a nested non-parametric MANOVA using the *BiodiversityR* package (60). We used *vegan* to perform PERMANOVAs (with 999 permutations) and principal coordinates analyses (PCoA) on the multiple dissimilarity matrices to determine the effect of age, clutch identity, tank, and sampling depth on microbiome composition. The function *betadipser* was used to calculate the distance between individuals and their age group centroids. We used *permutest* to determine if there were significant differences between mean distances to centroids between groups and Tukey’s Honest Significant Differences between groups to perform multiple group comparison for significant *permutest* results.

#### Effect of clutch over development

We then investigated whether the effect of clutch varied across development (i.e. whether eggs with shared parentage remained more similar than eggs without shared parentage across development time). To address this question, we used a Bayesian regression model with the R package *brms* (61–63) to model the effect of age and shared parentage (coded as 0 or 1 for different clutch or same clutch, respectively) on pairwise similarity (1-Jaccard Index) between eggs. A Bayesian regression model was ideal to investigate this relationship because it allowed us to control for non-independence in our dataset arising from pairwise measurements and uneven sampling of eggs derived from each stock tank (64). Since Jaccard similarity is bounded by 0 and 1, the models were fit to a beta distribution. Cross-tank pairs (A-B) were excluded from the model, because breeding pairs of fish were always from the same tank. The intercept and slope of this model varied by a multi-membership random effect capturing both eggs in each pairwise comparison (1 | mm[Individual_X_, Individual_Y_]), and by a random effect capturing clutch identity of the individuals in the pair. We tested the interaction effect between shared parentage and age pair (whether the pairwise comparison is between a 0 dpf and 0 dpf egg, a 0 dpf and 1 dpf egg, for all possible combinations), to determine whether the effect of shared parentage on the microbiome differs among development stages. We compared posterior means of shared clutch effect between different ages in the model using the “hypothesis” function in *brms*. The package *tidybayes* (65) was used to extract the predicted values of pairwise similarity from the model and calculate the relative effect of shared clutch on Jaccard similarity among eggs of different ages. For Bayesian regression models, if 95% credible intervals (CI) of the posterior distribution do not include 0, the term in question is considered to have a significant effect on the dependent variable.

#### Differential abundance analysis

We identified bacterial families and genera that were differentially abundant across development stage using a zero-inflated negative binomial regression model implemented in the R package *brms*. These models are ideal for microbiome count data, particularly when the data are over-dispersed and sparse (66,67). Each taxon was tested individually to determine whether the qPCR-scaled abundance of said taxon increased or decreased with egg age. A second model included read depth (scaled from 0 to mean abundance of all phyla, families, or genera) as a variable. Both models allowed the intercept to vary with respect to clutch. Prior to model fitting, the relative abundance of each ASV was scaled using 16S copy number calculated via qPCR (absolute abundance, Supplemental Methods). Sparse ASVs were removed based on read count abundance and prevalence across egg samples (ie. those ASVs not seen at least once in 5% of samples). All taxon models were run on 4 chains with 4000 iterations. Taxon models with over 100 divergent transitions (out of 8000) and R-hat values greater than 1.05 were discarded.

Posterior means were divided by the breadth of the credible intervals to calculate a weighted mean and account for uncertainty in the means. The package *ggtree* was used to plot the estimates for each taxon alongside its’ tip on a phylogenetic tree. To examine whether taxonomic class is associated with differentially abundant genera (i.e. whether genera within each class are associated with increasing abundance, decreasing abundance, or insignificant changes in abundance), we used a Chi-square test and plotted the Pearson residuals with the package *corrplot* (68).

## Results

We collected approximately 450 eggs from tank A, split across a total of 9 dishes, with approximately 50 eggs stored per dish. Parents from tank A laid fewer eggs than parents from tank B (9 dishes of approximately 50-75 eggs vs. 16 dishes). The only dish from tank A clutch 3 parents (A3) had fungal growth at 2 dpf, so these 2 dpf eggs were not collected. We collected approximately 800 eggs from tank B, split across 16 dishes. Of those dishes, we sampled eggs from 5 dishes collected from tank A, and 8 dishes collected from tank B. Five eggs per dish were sampled at each timepoint. At 1 day post fertilization, we removed 39 dead/nonviable eggs from tank A dishes and 14 from tank B dishes. Due to fungal growth in the clutch A3 dish at 2 days post fertilization, so we did not collect 2 dpf eggs from this dish (Supplemental Table 2). We collected a total of 124 eggs across the three ages and 8 clutches for subsequent microbial DNA extraction and sequencing.

After filtering using *dada2* and *decontam*, 27 samples had fewer than 2500 reads and were removed. For the subsequent analyses, we used 97 total eggs across 8 clutches and 3 development stages. Tank B parents produced more eggs than tank A, and after filtering, we had microbiome data from more than double the number of tank B eggs than tank A (67 eggs, compared to 30 eggs).

Across all ages, the egg microbiome was dominated by *Proteobacteria*. Among Proteobacteria, the most abundant and prevalent genera were *Rheinheimera*, *Pseudomonas*, and *Aeromonas* (Supplemental Figure 1A, Supplemental Figure 2). We also found that bacterial load increased significantly during development; log-transformed mean bacterial load had a significant linear relationship with age and increased between ages 0 and 2 dpf (Estimate = Estimate = 0.71, p = 0.00037; Supplemental Figure 1B).

### Phylogenetic diversity of the microbiome decreases during development

We measured alpha diversity using ASV richness (richness), Shannon Index (Shannon’s), Pielou’s evenness and Faith’s Phylogenetic diversity (PD) of the egg microbiome at fertilization (0 dpf), at 1 dpf, and immediately before hatching (2 dpf) to examine how microbiome diversity changes on zebrafish eggs. There were no significant differences in Richness, Shannon Index or evenness with respect to age (ASV Richness: F = 0.943, p = 0.393; Shannon: F = 0.879, p = 0.419; Evenness: F = 2.042, p = 0.136). However, there was a significant negative linear relationship between PD and age and a significant, but smaller positive quadratic relationship between PD and age (Figure 2A, Table 1). These results suggest that PD decreases exponentially shortly after fertilization. Post-hoc Tukey tests revealed that there were significant differences in PD between 0 dpf eggs and 1 dpf eggs (Estimate = 4.74, p_holm_ < 0.0001) and 2 dpf eggs (Estimate = 5.95, p_holm_ < 0.0001), but not between 1 dpf and 2 dpf eggs (Estimate = 1.21, p_holm_ = 0.121). These results indicate that a significant decrease in PD occurs within the first 24 hours post fertilization but does not decrease significantly after 24 hours. Additionally, the standardized effect size of PD (i.e. PD adjusted for ASV richness of sample) was significantly higher than expected for 0 dpf eggs (V = 379, p = 0.00011) and lower than expected for 1 and 2 dpf eggs (1 dpf: V = 56, p < 0.0001; 2 dpf: V = 0, p < 0.0001), indicating that the differences in PD across age groups were not the result of differences in richness.

**Figure 2:**
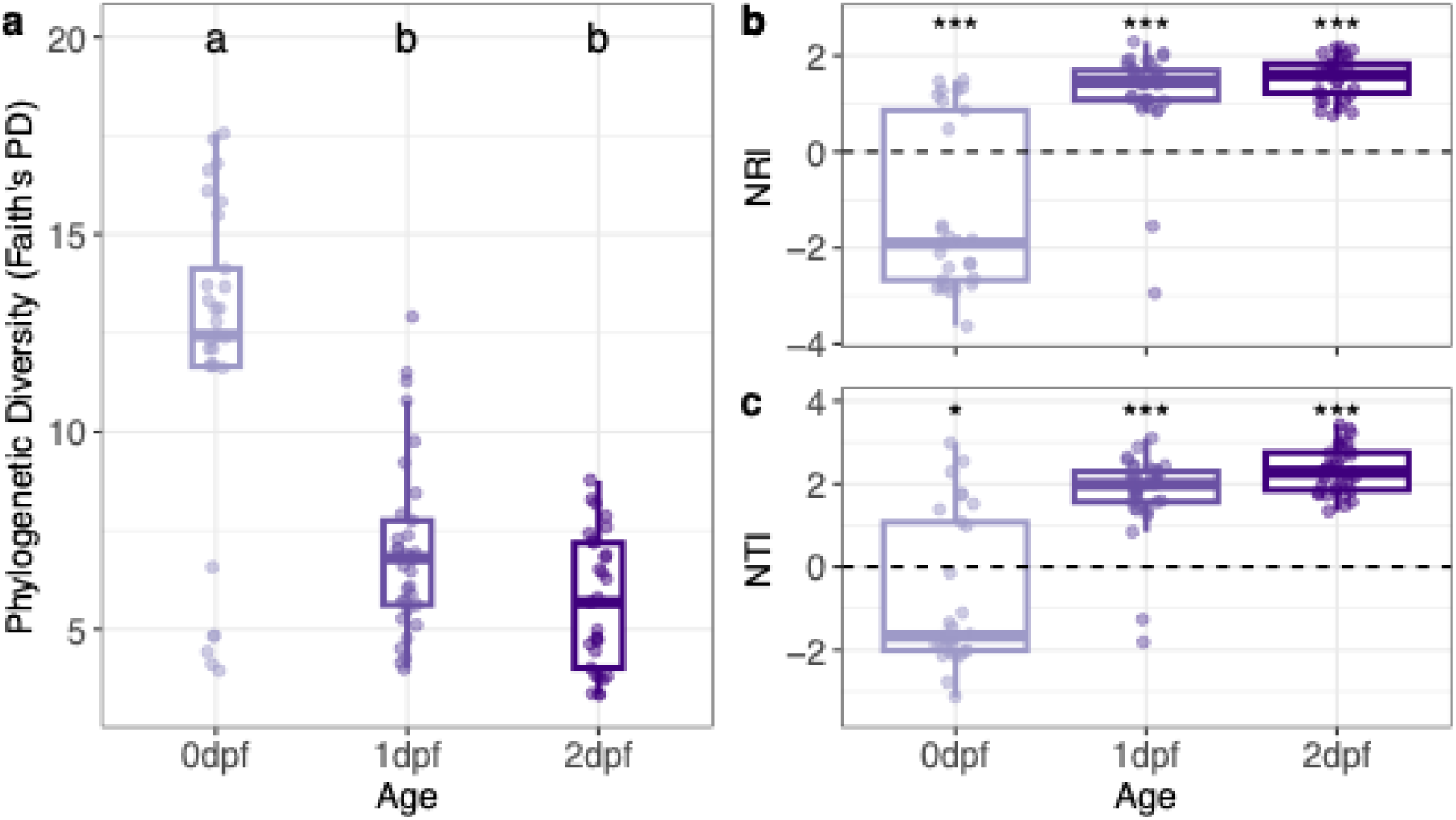
(A) Faith’s PD of eggs with respect to age (dpf). Means not sharing any letter are statistically significant according to multiple comparison tests with Tukey contrasts and p-value ≤ 0.05. P-values were adjusted using the Holm method. The NRI (b) and NTI (c) of egg microbiomes with respect to age (dpf). Asterisks denote means that are significantly less than 0 (0 dpf) or greater than 0 (1 dpf and 2 dpf) as determined by a one-sample Wilcoxon signed rank test (*: p < 0.05, ***: p < 0.0001).

**Table 1:** Results of regression models for measures of alpha diversity as a function of age and scaled read depth.

| Variable | Faith's PD |  |  | ASV Richness |  |  |
| --- | --- | --- | --- | --- | --- | --- |
|  | Beta | 95% CI | P | Beta | 95% CI | P |
| Age |  |  |  |  |  |  |
| Age | -4.2 | -5.2, -3.3 | **<0.001** | -2.3 | -6.2, 1.6 | 0.3 |
| Age<sup>2</sup> | 1.4 | 0.57, 2.3 | **0.001** | 0.96 | -2.6, 4.5 | 0.6 |
| Scaled Read Depth | 1.7 | -2.3, 5.7 | 0.4 | 17 | -0.12, 34 | 0.052 |
| Abbreviation: CI = Confidence Interval |  |  |  |  |  |  |

Since these patterns in PD could be due to environmental filtering (i.e. “sorting” of specific lineages by the environment of the developing egg), we calculated the net relatedness index (NRI) and nearest taxon index (NTI) for each microbiome, using the R package *picante*. If mean NRI and NTI are not significantly different from 0, this is an indication of random community assembly with respect to phylogeny. Positive NRI and NTI are indicative of phylogenetic clustering (a signal of environmental filtering), and negative values are indicative of overdispersion (a signal of competition). The mean NRI and NTI of the communities within each age group were significantly different from 0 (Figure 2B, Figure 2C) so the microbiomes were not randomly assembled with respect to phylogeny. Both NRI and NTI of 0 dpf eggs were less than 0 (NRI: V = 45, p < 0.0001; NTI: V = 126, p = 0.024). At 1 dpf and 2 dpf, NRI and NTI were significantly greater than 0 (NRI_1dpf_: V = 609, p < 0.0001; NTI_1dpf_: V = 649, p < 0.0001; NRI_2dpf_: V = 528, p < 0.0001; NTI_2dpf_: V = 528, p < 0.0001) suggesting a shift from a competitive to filtering processes in the first days of zebrafish microbiome development.

### Parentage and age explain variation in microbiome composition

Both parentage and development stage have been linked to compositional variation in the egg microbiome of several fish species (29,31,38). We investigated whether clutch identity (indicating parentage) and developmental age similarly influence the microbiome composition of zebrafish eggs, measured using Bray-Curtis, Jaccard, and Unifrac dissimilarity indices. Dish was not a significant source of variation between eggs when nested within clutch for any of the metrics of beta diversity (Bray-Curtis: F = 1.037, p = 0.327; Jaccard: F = 1.065, p = 0.356; Unifrac: F = 0.823, p = 0.485). Therefore, this term was left out of subsequent analyses.

There was no difference in mean dispersal of eggs (i.e. mean distance to centroid) between age groups when using Bray-Curtis, Unifrac and Jaccard as distance metrics (Bray: F = 0.604, p = 0.539; Jaccard: F = 0.786, p = 0.471; Unifrac: F = 1.865, p = 0.165). However, 1 dpf eggs were less dispersed in terms of Weighted Unifrac dissimilatory compared to 0 dpf eggs (p_Holm_ = 0.01) and 2 dpf eggs (p_Holm_ = 0.0003).

Egg microbiome composition varied by age, clutch identity (proxy for parental identity), and which stock tank the parents originated from (Figures 3A-C; Table 2). PERMANOVA tests found that ASV composition and phylogenetic composition varied significantly by age, parental stock tank, and clutch identity nested within tank (p < 0.001 for all metrics; Table 2). Scaled read depth also had a small, but significant effect on microbiome composition, but not phylogenetic composition (Bray: R^2^ = 0.014, p = 0.006; Jaccard: R^2^ = 0.015, p = 0.004; Unifrac: R^2^ = 0.009, p = 0.225).

**Figure 3:**
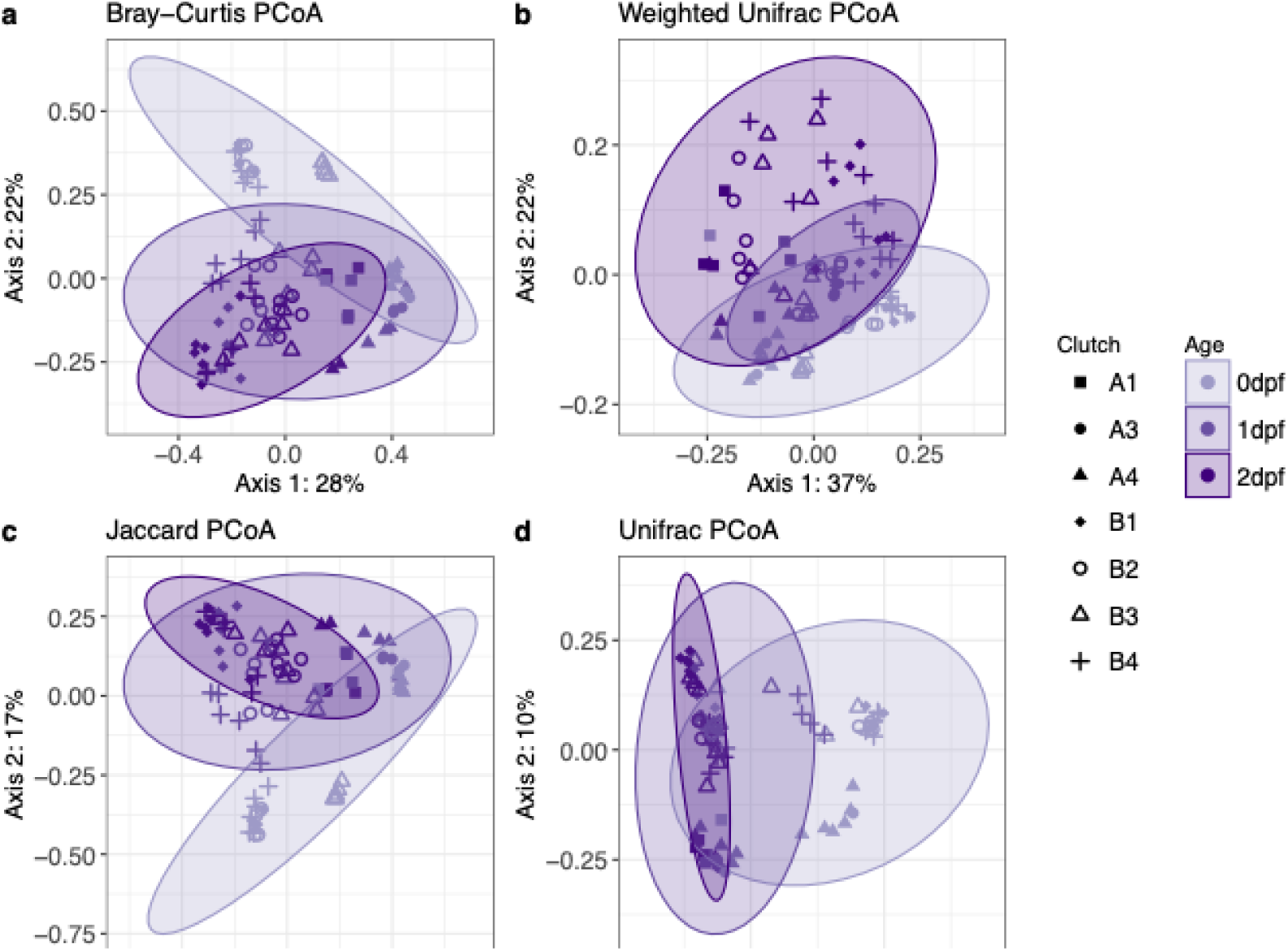
Composition of the zebrafish egg microbiome varies by age, clutch and parental stock tank. The principal coordinate plot (PCoA) show the (a) Bray-Curtis, (b) Weighted Unifrac, (c) Jaccard and (d) Unifrac dissimilarity between samples. Ellipses represent the 95% confidence interval around age groups assuming a multivariate t-distribution.

**Table 2:** PERMANOVA tests were performed on three difference distance metrics of the egg microbiome. Age, Tank and Clutch nested within Tank were provided as explanatory variables for similarities or differences between the microbiomes of individual eggs.

| Term | Bray-Curtis |  |  | Jaccard |  |  | Unifrac |  |  |
| --- | --- | --- | --- | --- | --- | --- | --- | --- | --- |
| | $R^2$ | F | P | $R^2$ | F | P | $R^2$ | F | P |
| Age | 0.216 | 22.166 | <0.001 | 0.177 | 14.183 | <0.001 | 0.236 | 16.698 | <0.001 |
| Tank | 0.215 | 44.142 | <0.001 | 0.150 | 24.051 | <0.001 | 0.086 | 12.236 | <0.001 |
| Tank:Clutch | 0.160 | 6.576 | <0.001 | 0.152 | 4.868 | <0.001 | 0.098 | 2.769 | <0.001 |
| Scaled Read Depth | 0.014 | 2.925 | 0.006 | 0.015 | 2.438 | 0.004 | 0.009 | 1.220 | 0.225 |
| Residual | 0.395 |  |  | 0.506 |  |  | 0.572 |  |  |
| Total | 1.000 |  |  | 1.000 |  |  | 1.000 |  |  |

### Age and clutch identity have varying effects on pairwise microbiome similarity

Overall, clutch identity and development stage significantly explained variation in microbiome composition between eggs. We next investigated whether the signal of clutch identity on microbiome composition changed over development. That is, does the signal of shared clutch status (i.e. shared parentage) on the egg microbiome decrease as the embryos approach hatching? We measured pairwise microbiome similarity using Jaccard Index, to calculate the proportion of shared ASVs between pairs of eggs and then used Bayesian regression modeling (Table 3) to estimate the effect of clutch on microbiome similarity on pairs of eggs across development (0 dpf-0 dpf, 1 dpf-1 dpf, 2 dpf-2 dpf for all possible age combinations).

**Table 3:** Posterior mean estimates and 95% quantile credible intervals for fixed effects from Bayesian regression model testing for the effect of shared clutch, age pairing, their interactions, stock Tank, and scaled read depth difference on Jaccard similarity. Bolded values indicate estimates significantly different from 0. ‘*’ denotes contrasts from hypothesis testing that have a posterior probability greater than 0.95.

| Term | Estimate (95 % CI) | Age pair x Shared Clutch (95% CI) | Hypothesis:<br>Shared > Different Clutch (95% CI) |
| --- | --- | --- | --- |
| **Intercept** | **-0.53 (-0.82, -0.26)** |  | 1.15 (1.00, 1.31) * |
| **Age pair** |  |  |  |
| **1dpf-1dpf** | 0.04 (-0.20, 0.27) | **-0.57 (-0.72, -0.42)** | 0.58 (0.43, 0.73) * |
| **2dpf-2dpf** | **-0.46 (-0.70, -0.23)** | **-0.60 (-0.75, -0.45)** | 0.55 (0.40, 0.70) * |
| **0dpf-1dpf** | **-0.63 (-0.76, -0.50)** | **-0.94 (-1.08, -0.81)** | 0.20 (0.07, 0.34) * |
| **1dpf-2dpf** | **-0.53 (-0.74, -0.33)** | **-0.81 (-0.94, -0.67)** | 0.34 (0.21, 0.48) * |
| 0dpf-2dpf | -1.14 (-1.27, -1.01) | -1.03 (-1.17, -0.89) | 0.12 (-0.02, 0.26) |
| Shared Clutch | 1.15 (0.96, 1.35) |  |  |
| Tank B | 0.20 (-0.06, 0.46) |  |  |
| Abbreviation: CI = Credible Interval |  |  |  |

Tank pair (A-A or B-B) was included as a population level effect. Pairs of eggs from tank B (B-B) were marginally more like each other than pairs of eggs from tank A-A (Jaccard: posterior mean [95% Credible Interval (CI)] = 0.20 [-0.06, 0.46]). In general, eggs originating from the same clutch had higher Jaccard similarity than eggs originating from different clutches (posterior mean [95% CI] = 1.15 [0.96, 1.35]). However, this effect of shared clutch varied depending on the ages of the eggs (Figure 4B). The posterior mean of the interaction term age pair * same clutch decreased during development (Table 3, Figure 4A), indicating that the effect size of shared clutch on microbiome similarity decreased as eggs aged.

**Figure 4:**
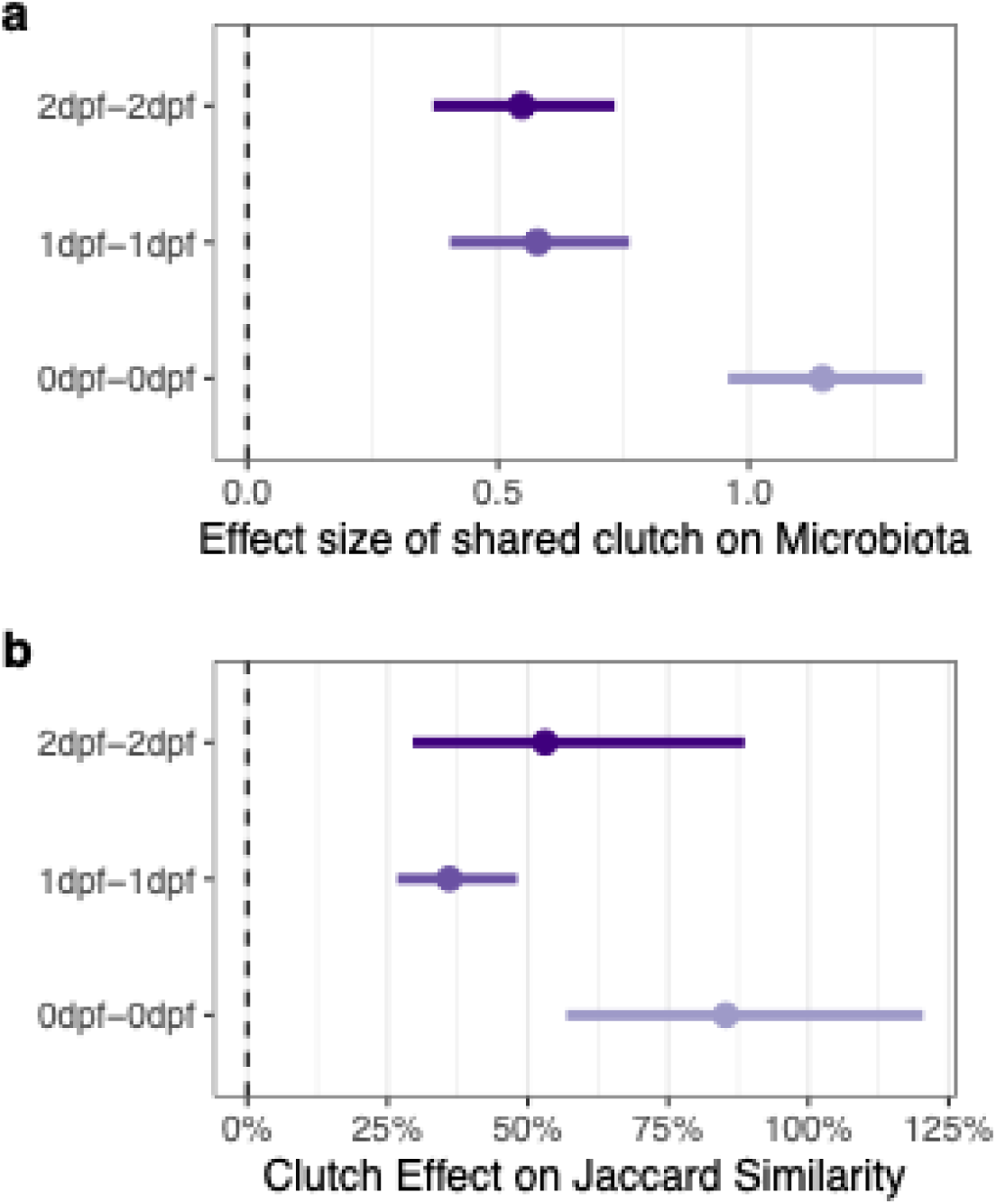
(A) Calculations from mean posterior draws of estimates from same clutch* age pair derived from BRM. (B) The clutch effect (i.e. the relative difference between pairwise similarity of eggs from the same clutches and different clutches) on Jaccard similarity with respect to the ages (dpf) of eggs in the pairwise comparison. The clutch effect was calculated from of the BRM predicted values and include all explanatory variables (Age Pair, Same Clutch, Tank pair, Clutch Pair).

We calculated the predicted values of Jaccard similarity for each egg according to the Bayesian regression model. At 0 dpf, eggs derived from the same clutch were 85.2% [95% CI: 56.7%, 120%] more similar to each other in terms of shared ASVs than eggs derived from different clutches. The effect of shared clutch on Jaccard similarity decreased to 36.0% [95% CI: 26.9%, 47.9%] for 1 dpf eggs, and increased to 53.0% [95% CI: 29.5%, 88.1%] for 2 dpf eggs.

### Genera belonging to Alphaproteobacteria are more abundant in 2 dpf eggs than 0 dpf eggs

After removing rare ASVs, the remaining 333 ASVs were aggregated to the genus level. ASVs unclassified at the genus level were removed. The resulting OTU table represented the absolute abundance of 156 genera. Thirty-three genera were discarded because the Bayesian regression models failed to converge (intercept and estimate Rhats < 1.05). The absolute abundance of 43 out of 123 bacterial genera significantly increased or decreased over egg development (Figure 5B). A Chi-squared test revealed that genera that are differentially more abundant or less abundant tend to be associated with higher level taxonomic groups (χ = 19.55, p = 0.033; Figure 5B). For instance, within the class Alphaproteobacteria, there were more genera that were increasing in abundance with age and fewer genera that were decreasing in abundance than would be expected, whereas Gammaproteobacteria and Bacteroidia exhibited the reverse pattern.

**Figure 5:**
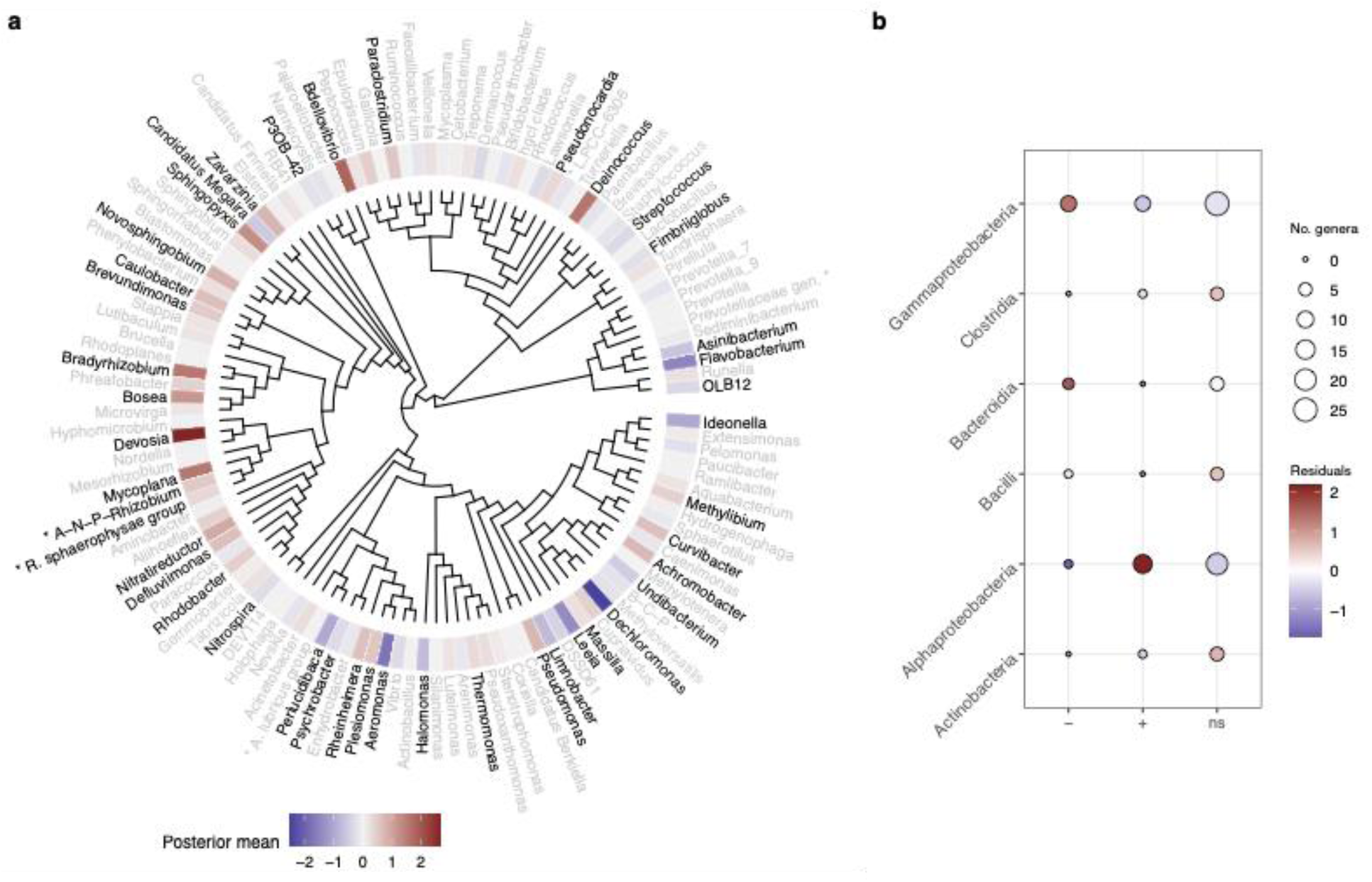
Cladogram (A) of genera with significant linear relationships with age. Bars at the tips of the tree represent the weighted posterior mean calculated from the regression models. Genera marked with asterisk (*) have abbreviated labels (A-N-P-Rhizobium = Allorhizobium-Neorhizobium-Pararhizobium-Rhizobium; B-C-P = Burkholderia-Caballeronia-Paraburkholderia; M-M = Methylobacterium-Methylorubrum; Prevotellaceae gen. = Prevotellaceae UCG-003; A. lubricus group = Agitococcus lubricus group. Correlation plot (B) of taxonomic classes and differentially abundant genera. Red represent positive associations, and blue represent negative associations. The size of the points represent the total number of genera within that class.

## Discussion

In this study, we address the effect of developmental stage and parental identity on microbiome composition of zebrafish eggs. These eggs were derived from wildtype adult zebrafish housed in the University of Oregon’s Huestis zebrafish facility. This study focuses on microbiome composition of the eggs during embryonic development, from 0 days post fertilization to 2 days post fertilization and uses 16S rRNA gene amplicon sequencing to survey the bacterial microbiome on the eggs. We found that the egg microbiome changes dramatically from fertilization to hatching. Parentage was also a significant driver of microbiome similarity.

We used 16S rRNA amplicon sequencing to survey the egg microbiome. This method has allowed us to track compositional differences in the egg microbiome due to age or parentage. However, 16S rRNA amplicon sequencing has limited resolution, especially for strain-level identification (69), and may not accurately represent the functional diversity present in the egg microbiomes. The utility and accessibility of 16S rRNA amplicon sequencing make it an acceptable tool to address the questions in this study, despite its limitations. Future zebrafish microbiome studies may incorporate functional profiling and metagenomic to determine strain-level diversity and determine the functions of the egg microbiome.

### Microbial transmission likely explains within clutch similarity

We found that parentage significantly explained variation in the egg microbiome across age groups. Since dish nested within clutch was not a significant source of variation, the similarity of eggs with shared parentage is not due to cohousing of the eggs. High similarity in the microbiomes of eggs with shared parentage may be the result of either transmission of microbes from parent to offspring, or similarities due to host genetics. Transmission and host genetics may explain why eggs within the same clutch have more similar microbiomes compared to eggs from different clutches. We propose that transmission may be a likelier explanation for microbiome similarity due to parentage.

Firstly, we observed a large effect of tank on microbiome composition. In all metrics of beta diversity, the stock tank of the parents left a significant signal on microbiome composition. These results suggest that housing effects may be passed on to subsequent generations of animals. While it is possible these tank differences could be confounded by the genetic relatedness between the adults, this is also unlikely since adult zebrafish in the UO Heustis Zebrafish Facility are only segregated by genotype, and not by parentage. Therefore, the similarity by tank is likely the result of transmission microbes between generations. Cohousing is known to affect the microbiomes of animals (70), and these results show that cohousing can also affect the microbiomes of subsequent generations. Future studies could address whether this tank effect on the microbiomes persists when clutches derived from separately housed parents are combined and cohoused.

Secondly, using Bayesian regression models, we observed the effect of shared parentage decreased as the eggs developed. If the microbiome similarity within clutches was solely due to similarities in host genetics, we would likely not observe a decay in clutch effect over development. Transmission of microbes from parent to offspring may be responsible for the high similarity between closely related zebrafish eggs, particularly shortly after fertilization. Other factors explaining the decay may be physiological changes over embryonic development, or the embryo’s production of anti-microbials (39). Future studies will incorporate 16S rRNA sequencing of parental tissues and environmental sources (in this case, facility water) and use source-sink inference to quantify the contributions of potential sources to the egg microbiome, and whether these contributions change over time.

### Age explains microbiome variation and compositional shifts

Age explained more variation than parentage on the microbiomes of zebrafish eggs across beta-diversity metrics (Table 2). Additionally, age correlated with a decrease in phylogenetic alpha diversity (Faith’s PD; Figure 2A). This result could be due to environmental filtering of the microbiome by the developing egg, such as chemical changes in the chorion (71) or the production of anti-microbials (39). Environmental filtering, aka species sorting, can lead to phylogenetic clustering, where members of a microbiome are more closely related than predicted by random microbiome assembly. While competition may be driving assembly at 0 dpf (Figure 2B-C), environmental filtering may drive assembly on 1 and 2 dpf eggs and lead to phylogenetic clustering in the microbiome. These patterns may arise due to decreasing bacterial abundance across all taxa. However, using qPCR to quantify 16S copy number, we found that overall bacterial load increased from 0 dpf to 2 dpf (Supplemental Figure 1B).

We also identified several differentially abundant genera across age groups, the majority belonging to the classes Gammaproteobacteria and Alphaproteobacteria. Members of these classes are prevalent and abundant in the gut microbiomes of both larval and adult zebrafish across facilities (72) and development stage (26). We found that 9 of 43 genera (21%) within class Gammaproteobacteria decreased in abundance over development. Notably, *Aeromonas*—a genus within Gammaproteobacteria that have been identified as prevalent and/or abundant members of zebrafish adult and larval gut microbiomes (26,66,72)—was present in the egg microbiomes, and tended to be less abundant in 2 dpf eggs than in 0 dpf eggs. *Halomonas* followed this same pattern. This genus may be an intestinal pathogen of zebrafish (73). Stephens et. al 2016 found *Halomonas* was more prevalent in larval and juvenile zebrafish intestines than in adults. *Halomonas* may be acquired by zebrafish through brine shrimp feeding and later transmitted to the eggs if it persists in the adult gut (74). Eight genera within Gammaproteobacteria (18%) increased in abundance. Among these were ASVs belonging to *Pseudomonas*, *Rheinheimera*, *Achromobacter*, *Plesiomonas* and *Curvibacter*. *Pseudomonas* spp., *Plesiomonas* spp. and *Curvibacter* spp. have been previously detected in larval zebrafish raised in aquaria (26,75). *Achromobacter* is a rare member of the zebrafish gut, but was found to be readily transformed in the zebrafish gut through plasmid conjugation (76). Both *Rheinheimera* and *Pseudomonas* were enriched in conventionalized zebrafish larvae and may impact the neurodevelopment, although the mechanism by which this occurs requires further investigation (6).

Fourteen of 35 genera (40%) within Alphaproteobacteria increased in abundance, notably *Devosia*, *Mycoplana, Bradyrhizobium, Bosea* and *Sphingopyxis.* Members of these genera have been isolated from diverse environments (e.g. soil, marine and fresh water, root nodules and contaminated environments) and are often aerobic and motile (77–82). *Devosia, Mycoplana,* and *Sphingopyxis* have also been identified as core members of the zebrafish larval microbiome in one study (83).

Additionally, the genus *Flavobacterium* (class Bacteroidia), in which there are several known fish pathogens (84–88), significantly decreased in abundance over development. These patterns may indicate that the egg chorion or embryo medium is an amicable habitat for aerobic bacteria that are adapted for aquatic environments, or the early larval gut, more so than bacteria commonly associated with adult zebrafish intestines.

Metagenomics or metabolic profiling of the microbiome could provide insight into why these bacteria increase of decrease in abundance. However, the association between increasing or decreasing abundance and bacterial class is congruent with our findings of increased phylogenetic clustering as the eggs age. The loss of bacteria that tend to be associated with the zebrafish guts indicates that the eggs are not an amicable environment for typical zebrafish commensals (and pathogens) and that these bacteria must be acquired through other sources, such as the facility water system or through feeding.

Our results indicate that age and shared parentage have a significant and substantial impact on egg microbiome composition. The microbiome associated with fish eggs has been shown to change over development in other fish species with longer embryonic stages than zebrafish, such as brown trout and Atlantic salmon (29,37). We found that the egg microbiome of zebrafish changes rapidly, both in phylogenetic diversity and composition. The rapid changes in the microbiome may be due to the environmental conditions in the embryo media, or species sorting by the egg. Immunocompetent factors are present inside the eggs of several fish species (89,90), including zebrafish (36,39). However, it is unknown whether these factors can inhibit growth of microbes on the outside of the egg, since the chorion of fish eggs is likely impermeable to large organic molecules (91). Future research could delve into how physiological changes of the host and chorion during embryonic development might recruit or inhibit growth and attachment of bacteria to the chorion.

### Future directions

In fish species such as Atlantic salmon, the egg microbiome can provide defenses against pathogens (87,92). Future studies could use a combination of culturing, metagenomics, functional predictions from phylogeny, and microbiome manipulation to determine which taxa and/or functions are beneficial for developing zebrafish eggs. Furthermore, it will be important to establish whether these members of the egg microbiome are transient or if they can colonize the larval gut post-hatching. If the egg microbiome is an important source of microbes to the larval gut, there could be consequences later in development, revealing an important life stage during which microbiomes could be manipulated to improve survival of larval and juvenile fish.

Combating pathogens is an important consideration for aquaculture, where detrimental host-microbe interactions can lead to high mortality, especially among larval and juvenile fish (93). Microbiome manipulation and probiotics are potential alternatives to antibiotic treatment; however research on the effectiveness of probiotics has shown mixed success (94–96). Understanding the processes of microbiome assembly in different fish species and under different rearing conditions is vital to the development of efficient methods for microbiome manipulation to benefit host health. This study is a first step toward understanding the processes that shape the early-life microbiome assembly of zebrafish reared in an aquaculture facility.

Here we characterize the egg microbiome of zebrafish and investigate the major drivers of variation. Early-life assembly of microbiomes can have health and developmental consequences across the life span of a host; however, little is known about the drivers of early-life microbiome assembly in the model vertebrate, zebrafish. In zebrafish-microbe interaction studies, eggs are often sterilized to render them (and the larvae that subsequently develop) axenic. Egg sterilization is also a common practice in aquaculture, where a major concern is reducing the load of pathogens. However, these practices may eliminate potentially beneficial microbes that could be acquired from the parents or the environment in which the eggs were spawned. These bacteria may also persist on the eggs and be later acquired by the larvae.

Therefore, understanding the drivers of microbiome assembly on fish eggs may reveal previously unknown commensal microbes, and early members of the larval gut microbiome, which has been shown to be integral for larval health and development in zebrafish (1,2,6).

## Data Availability

16S rRNA gene amplicon sequences were deposited in a BioProject under the accession number PRJNA1266191. All code used in data analysis for this manuscript has been deposited on GitHub: https://github.com/biocsmith/ZFISH_microbiome_dev.

